# Commitment to cytokinetic furrowing requires the coordinate activity of microtubules and Plk1

**DOI:** 10.1101/2024.09.16.612913

**Authors:** Charles A. Day, Alyssa Langfald, Tana Lukes, Hanna Middlebrook, Kevin T. Vaughan, David Daniels, Edward H. Hinchcliffe

**Affiliations:** Cellular Dynamics Section, The Hormel Institute, University of Minnesota Austin, MN 55912 USA; Department of Biological Sciences, University of Notre Dame, Notre Dame IN 46556 USA; Dept. of Pediatrics and Neurosurgery, Mayo Clinic, Rochester, MN 55905 USA; Masonic Comprehensive Cancer Center, University of Minnesota, Minneapolis, MN 55455 USA

**Author notes:** Correspondence should be addressed to EHH.

**Keywords:** Astral microtubules, Aurora B kinase, anillin, central spindle, cytokinesis, furrowing, microtubule, midzone, mitosis, Polo-like kinase 1, RhoA

## Abstract

At anaphase, spindle microtubules (MTs) position the cleavage furrow and trigger actomyosin assembly by localizing the small GTPase RhoA and the scaffolding protein anillin to a narrow band along the equatorial cortex [1–6]. Using vertebrate somatic cells we examined the temporal control of furrow assembly. Although its positioning commences at anaphase onset, furrow maturation is not complete until ∼10-11 min later. The maintenance of the RhoA/anillin scaffold initially requires continuous signaling from the spindle; loss of either MTs or polo-like kinase 1 (Plk1) activity prevents proper RhoA/anillin localization to the equator, thereby disrupting furrowing. However, we find that at ∼6 min post-anaphase, the cortex becomes “committed to furrowing”; loss of either MTs or Plk1 after this stage does not prevent eventual furrowing, even though at this point the contractile apparatus has not fully matured. Also at this stage, the RhoA/anillin scaffold at the equator becomes permanent. Surprisingly, concurrent loss of both MTs and Plk1 activity following the “commitment to furrowing” stage results in persistent, asymmetric “half-furrows”, with only one cortical hemisphere retaining RhoA/anillin, and undergoing ingression. This phenotype is reminiscent of asymmetric furrows caused by a physical block between spindle and cortex [7–9], or by acentric spindle positioning [10–12]. The formation of these persistent “half-furrows” suggests a potential feedback mechanism between the spindle and the cortex that maintains cortical competency along the presumptive equatorial region prior to the “commitment to furrowing” stage of cytokinesis, thereby ensuring the eventual ingression of a symmetric cleavage furrow.

## Results and Discussion

At the end of mitosis, the cytokinetic furrow forms along the cell equator, and is both initiated and positioned by the spindle MTs [13–18]. These MTs activate a band of RhoA at the equator via kinesin-6 transport of the RhoA activating/inactivating enzymes MgcRacGAP and Ect2 [4, 19]. Coordinate with RhoA activation in the presumptive furrow is the localization of the scaffolding protein anillin, which links together RhoA, actin, myosin and the plasma membrane [5, 6, 20, 21]. MTs also transport Aurora B (Aur B) kinase to the equatorial cortex, where it may activate kinesin 6 [18, 22]. The stimulation and deposition of these furrowing activities, particularly active RhoA, occurs immediately after anaphase onset [2]. Early in cytokinesis, the cortex remains plastic, as the localized band of activated RhoA moves in response to spindle re-positioning. This suggests that the contractile machinery is not fully assembled [3]. These data indicate that the positioning of the furrow can be manipulated up to a “point of no return”, after which contractility proceeds [2, 3, 16]. Where this “point of no return” lies, relative to anaphase onset, remains unknown.

### Furrow Assembly in MT-free Cells

To examine the timing of furrow assembly and ingression in response to MT-based cleavage signals, we utilized a single-cell based MT depolymerization/re-growth assay based on vertebrate somatic (BSC1) cells expressing α tubulin-GFP [23, 24]. Beginning in metaphase, cells were imaged by spinning disk confocal microscopy, and the time from anaphase onset to the onset to cortical furrowing (judged by a 20% decrease in furrow width) was measured. In control cells, this is 10 +/- 1.6 minutes (Figure S1A; n = 16). At defined intervals following anaphase onset, the cell of interest was chilled to 4 °C for 20 min to depolymerize the spindle MTs (Figure 1, also see Figure S1). Chilled cells were then returned to the pre-warmed microscope and MT re-assembly was imaged. Initially, no MTs were visible, though the spindle poles were often seen as a pair of bright fluorescent dots (Figure S1D). Within two minutes, re-warmed cells assembled both astral MTs and a spindle midzone, and anaphase proceeded. As a control, direct fixation of cells following chilling revealed that all of the MTs had disassembled, including those of the midzone, as evidenced by the lack of PRC1 labeling (Figure S1B). However, chilling cells did not disrupt the localization of cleavage furrow components RhoA and anillin to the equatorial cortex (Figure S1C).

**Figure 1.**
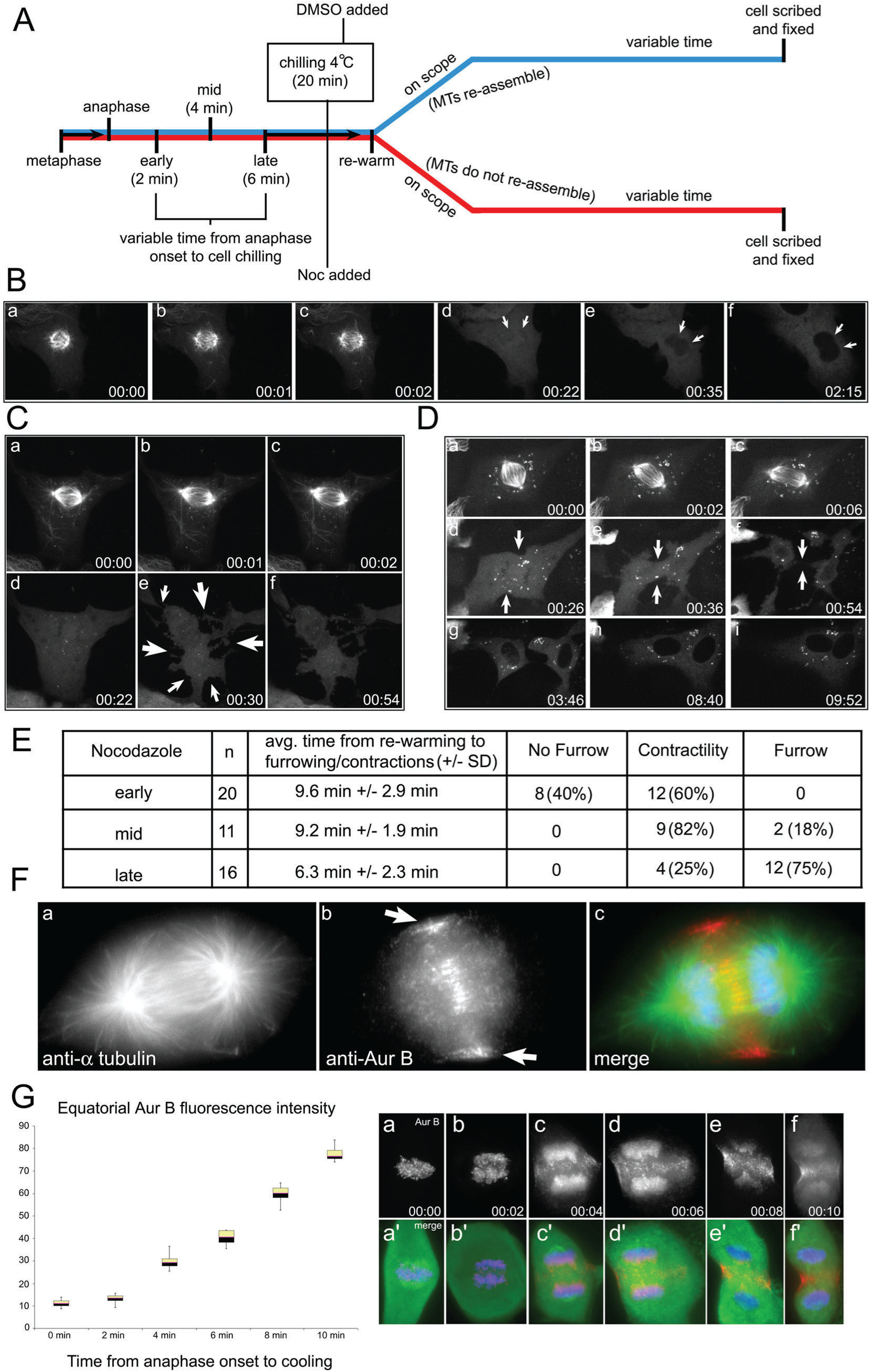
Suppressing MT re-assembly after chilling results in cytokinetic defects during re-warming. **A.** Schematic of cold-based spindle disassembly. Blue timeline depicts the MT re-assembly route (control), and the red timeline depicts the experimental route (MT re-assembly is inhibited). Cells were followed as they enter anaphase, and then chilled/re-warmed in the presence of 33 μM Nocodazole. There are three resulting phenotypes. **B.** Early anaphase, no furrow formation. There is not furrow, and no cortical contractility. **C.** Early anaphase, non-specific cortical contractility. The cell forms multiple cortical ingressions (arrows). These eventually retract. **D.** Late anaphase, a single broadened furrow forms. The cell forms a single furrow along the equator that constricts the cell into two (**f**). These furrows eventually retract over a period of hours, and the cells fuse. **E.** Average time from re-warming to the induction of a furrow and percentage of each phenotype for cells chilled in early, mid, or late anaphase. Hrs:Min. **F. a-a”.** Control cell, fixed in anaphase, showing the lateral equatorial localization of Aur B (arrows) along with the midzone localization of Aur B bound to the CPC. **G.** Quantification of fluorescence intensity of Aur B at the equatorial cortex in chilled anaphase cells. Box and whiskers plot; the box represents the 25^th^ – 75^th^ percentile, the central line is the median value, and the error bars represent the full range of intensity values for each time point. N = 6 cells measured for each time point. **a-f.** Time-course of Aur B re-distribution to the equatorial cortex. As cells progress through anaphase, the centromeric localization of Aur B is lost between 0 and 4 minutes post-anaphase. Over 4 to 10 minutes post-anaphase there is an increase in the equatorial cortical localization of Aur B (**c-f**). **a’-f’.** Overlay of Aur B, anti-α tubulin and DAPI. Bar = 10 μm.

For these and subsequent experiments, cells were chilled at three points during anaphase: *early* (2 minutes after anaphase onset), *mid* (4 minutes post-anaphase), or *late* (6 minutes post-anaphase). In all cases (n=31), a single cleavage furrow assembled that was restricted to the equatorial region of the cell. The average time from re-warming to furrow initiation was dependant upon how far anaphase had progressed prior to chilling: for *early* anaphase cells it is 8.0 +/- 1.2 min., for *mid*-anaphase cells, 9.7 +/- 2.1 min., and for *late* anaphase cells it is 6.5 +/- 1.7 min. (Figure S1E).

To examine the effects of completely blocking spindle MT re-assembly during cell re-warming, cells were chilled to 4 °C in the presence of 30 μM Nocodazole (Noc) in *early*, *mid*, and *late* anaphase respectively (Figure 1). These cells were then re-warmed, and cytokinesis was imaged. Under these conditions, all spindle MTs were disassembled, and did not reassemble upon warming. Figure 1 shows a schematic for our disassembly-reassembly experiments, and the resulting outcomes for cytokinesis in each class.

In *early* anaphase cells, inhibiting MT re-assembly resulted in either a complete lack of cortical contractility (40%; Figure 1B), or extensive random cortical contractility, with no bias towards the cell’s equatorial region (60%; Figure 1C). In those cells that lack cortical contractility, the nuclei reformed in a common cytoplasm, and the cells exited mitosis as single daughters (Figure 2B, d-f – arrows). Same cell live-fixed analysis revealed that these cells lack a centrally localized band of RhoA (N = 4) and anillin (N = 4; Figure S2).

**Figure 2.**
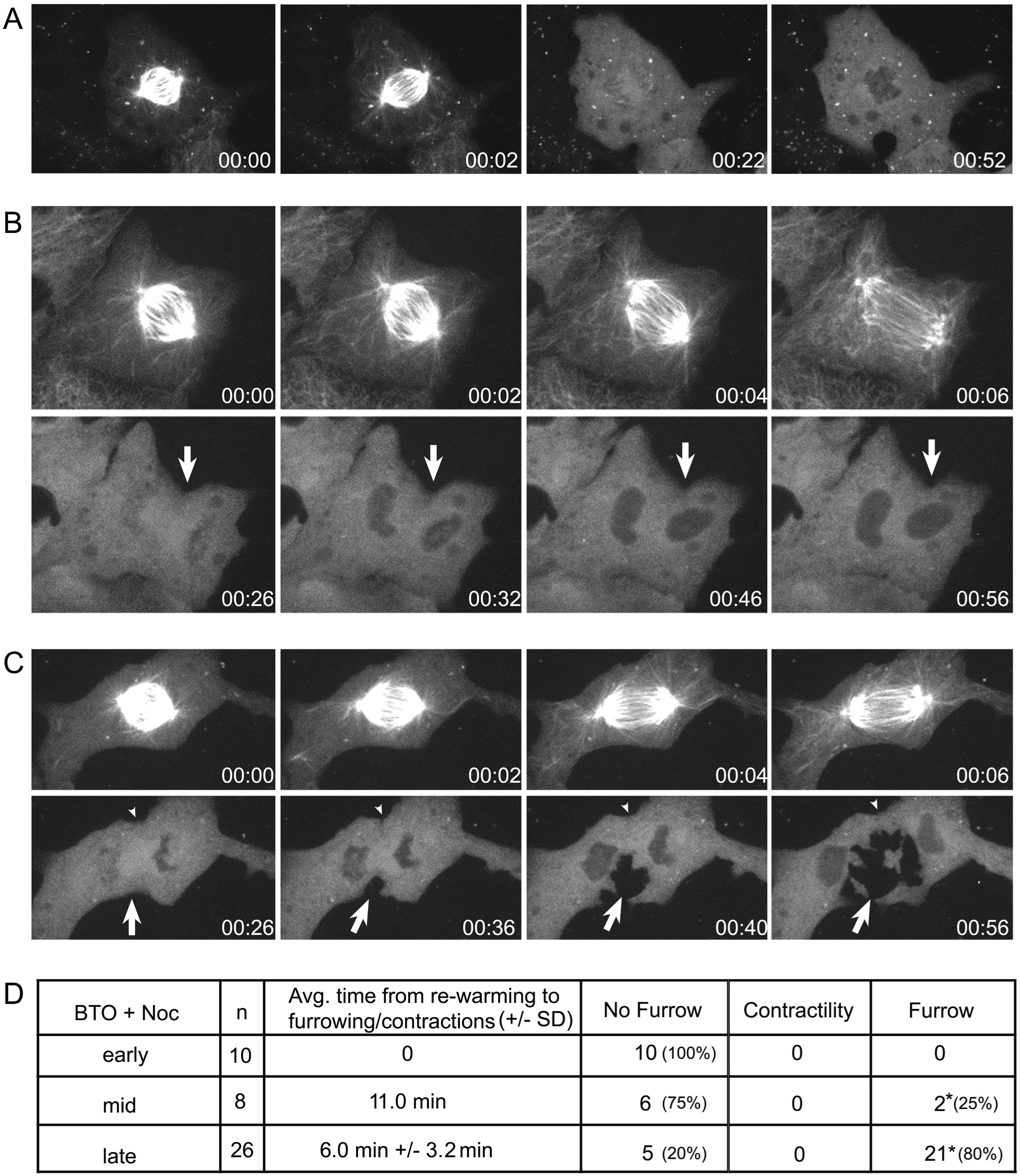
Plk1 activity regulates cortical contractility in the absence of microtubules. Cells chilled in early (**A**) or mid-anaphase (**B**) and treated with BTO-1 + Noc. Upon re-warming, these cells fail to re-form spindle MTs, and do not undergo cortical contractility nor assemble a furrow. **C.** A cell chilled in late anaphase (6 min) and then BTO-1 + Noc added. As this cell is re-warmed (26 min), an asymmetric cleavage furrow forms on one surface of the cell (large arrow). This “half-furrow” ingresses and the cell elongates. Note that the cortex lying opposite of the “half-furrow” remains quiescent (small arrowhead). **D.** Number of cells and furrowing phenotype for early, mid- and late chilled cells treated with BTO-1 + Noc. Note that in this case the furrow formation in the late anaphase category (**\***) refers to “half-furrows” only. Time is hrs:min. Bar = 10 μm.

Chilling cells in the presence of Noc beginning in *mid* anaphase (Figure 1E) resulted in the majority exhibiting the random cortical contractility phenotype (82%), with a few managing to direct the formation of a prominent furrow (18%). In the case of random cortical contractility, post-anaphase cells assembled multiple regions of cortical ingression [see 13], but lacked a prominent central cleavage furrow (Figure 1C, arrows in panel e). Same cell immuno-analysis suggests that although cortical RhoA (N = 3), anillin (N = 4) and actin/myosin (N = 3) are present in these cells, there is no concentration at a defined furrow (Figure S3).

When cells were allowed to progress into *late* anaphase (6 minutes), 25% underwent non-specific contractility without a furrow, whereas 75% formed a prominent centralized furrow. Those cells that managed to form a furrow cleaved into two daughters connected by a thin bridge of cytoplasm (Figure 1D). However, because these cells lacked a midbody (caused by a lack of MTs), abscission could not take place [25–27], and after a period of several hours the two daughters fused into a single binucleate cell (Figure 1D). Interestingly, same cell immuno-analysis of these late anaphase cells revealed that the distribution of RhoA (N = 3) and anillin (N = 3) extended along a broadened furrow, even after the MTs were depolymerized (Figure S2C, D). Thus, at this point, the cells have received signals sufficient to promote focused furrowing at the equator, and the continued presence of MTs is not required.

Previous work suggests that as time in anaphase progresses, there is first an induction of global cortical contractility, and then the refinement of cortical ingression into the equatorial region of the cells, building a focused furrow [28]. To test this idea, we measured the deposition of Aur B kinase at the equatorial cortex as a marker for astral MT delivery (Figure 1F). At anaphase, Aur B exists in two sub-cellular populations: (i) as a component of the Chromosome Passenger Complex (CPC), which transits from the centromeres to the central spindle at anaphase onset, and (ii) a cytoplasmic population, which is delivered to the equatorial cortex via astral MTs [22]. To measure cortical Aur B, we followed individual cells as they transited anaphase, chilled them at 2 min intervals, then fixed them directly for IF analysis. This revealed a 4-fold increase in Aur B localization to the cortex during the period from 2 to 6 minutes post-anaphase (Figure 1G). To confirm that this cortical localization was dependent solely on astral MTs, early anaphase cells were chilled and a low dose of Noc added, which suppressed astral MT but not midzone MTs re-growth upon re-warming (Figure S4). In such cells, there is clear localization of midzone Aur B (the CPC population), but not cortical Aur B (Figure S4B).

These data suggest that beginning early in anaphase there is a deposition by the MTs of signals to the equatorial cortex that induce furrowing. These signals are continuously required for a period of ∼6 min following anaphase onset. After this time, the cortex “commits” and is capable of furrowing without the continued presence of any MTs, astral or midzone. Furthermore, during early anaphase, loss of astral MTs prevents the localization of cleavage factors (such as Aur B) to the equatorial cortex (Figure 1G). If anaphase is allowed to progress for longer than 2 minutes, then there is a progressive increase in the amount of furrowing activities brought to the cortex by astral MTs, with a commensurate increase in furrow formation and focusing.

These results support the idea that the complete loss of spindle MTs beginning in very early anaphase prevents the formation of a defined furrow, as these cells lack an equatorial zone of RhoA necessary to continuously induce furrowing [16, 29]. These cells either lack furrowing, or lack the ability to focus the furrow to a single equatorial band. As cells progress into mid-anaphase, loss of MTs results in an increase in the induction of global cortical contractility, but a reduced ability to focus the cortical ingressions into a distinct furrow. However, in late anaphase (∼6 min), it appears that sufficient signals have reached the cortex to support a radially symmetric furrow, and at this point in anaphase furrowing does not require any further signals from the astral MTs or spindle midzone in order to commit to cleavage. Thus, we propose that this time represents a “commitment to furrowing” phase during cytokinesis.

### MTs and Polo-like kinase 1 Activity Together are Required to Propagate a Radially Symmetrical Cleavage Furrow

Plk1 is a key regulator of mitosis, and is essential for spindle MT organization, kinetochore function, and cytokinesis [24, 30–32]. To examine the role of Plk1 during MT-free cytokinesis, we used cold-based MT depolymerization, along with Noc treatment to prevent MT re-growth, and also added the small molecule inhibitor BTO-1 to block Plk1 activity [24]. We chilled cells in *early*-, *mid*-, and *late*-anaphase, and examined their ability to initiate and ingress a cleavage furrow following re-warming (Figure 2). When both MTs and Plk1 activity were inhibited in either *early* or *mid*-stage cells, we did not observe furrowing, nor non-specific cortical contractility (Figure 2A). However, cells allowed to progress into cytokinesis for 6 min before chilling, followed by MTs and Plk1 inhibition generated a persistent, asymmetric “half-furrow” upon re-warming (Figure 2C). This “half-furrow” ingressed towards the cell center (Figure 2C, arrow), while the equator of the opposite hemisphere remained quiescent (Figure 2C, arrowhead). Half-furrows were observed in 80% (21/26) of cells chilled in late anaphase with Noc and BTO-1, whereas 20% of these cells did not undergo any furrowing (Figure 2B).

We next determined whether asymmetric furrowing represented (i) a lack of furrowing component deposition along the circumference of the cortex, or (ii) a differential activation of these proteins at regions that underwent contractility. First, we examined the localization and distribution of the furrow components Aur B, RhoA, anillin, MyoII and F-actin in cells that assembled “half-furrows” (Figure 3). Individual cells were followed as they progressed into anaphase for 6 min, then chilled to 4 °C for 20 min in the presence of Noc and Plk1 inhibitors, then re-warmed on the spinning disk confocal microscope. After the “half-furrow” had formed, the position of the cell was marked with a diamond scribe, and the coverslip fixed and labeled with the appropriate antibody. Aur B (N = 3), RhoA (N = 6), MyoII (N = 6) and anillin (N = 4) all localized along the surface of the “half-furrow”, but were absent from the opposite, non-contractile surface of the equatorial cortex (Figure 3). Interestingly, F-actin (N = 3) was found along the entire cortical surface, with no obvious bias to the “half-furrow” (Figure 3D).

**Figure 3.**
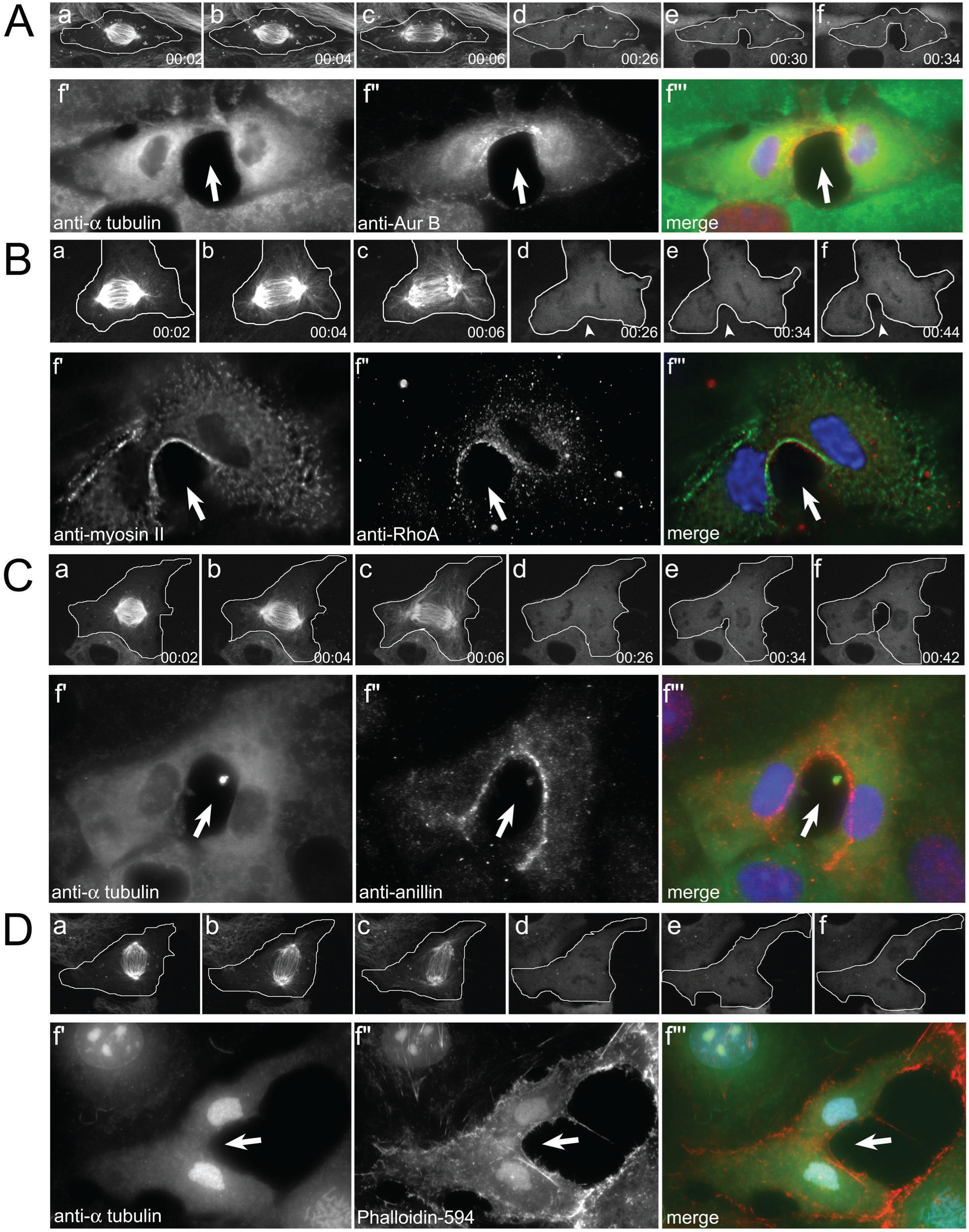
“Half-furrows” preferentially accumulate Aur B, Rho A, MyoII, anillin, but not F-actin. Same cell live/fixed images of cells with half-furrows induced by inhibiting Plk1 and MTs. **A.** Aur B localizes along the inner surface of the asymmetric furrow (f”, arrow). **B.** Both Myosin II and Rho A localize to the half-furrow (arrows). **C.** Anillin localizes along the ingressing surface, but not on the opposite side. **D.** F-actin, labeled with fluorescent phalloidin, is localized along the entire cell cortex, and is not concentrated on the surface of the ingressing furrow. Time = hrs:min, bar = 10 μm.

Next we compared the effects of inhibiting Plk1 activity alone during MT re-warming (Figure S5). Cells chilled in either *early*-anaphase or *mid*-anaphase and treated with BTO-1 reassembled prominent astral MT arrays, but these could not induce furrowing or complete cytokinesis (Figure S5). In addition, Plk1 inhibition during early anaphase did not inhibit the localization of anillin (N = 4) to an equatorial band, but did prevent equatorial accumulation of RhoA (N = 6; Figure S5A). This is consistent with work in HeLa cells, where blocking Plk1 activity in early anaphase prevented RhoA accumulation at the equatorial cortex [30].

Not surprisingly, blocking Plk1 activity in cells chilled/re-warmed in early anaphase inhibited the assembly of a spindle midzone, as judged by labeling with anti-MKLP1 (N = 4) and anti-PRC1 (N = 4; Figure S6). However, the most striking feature of cells re-warmed in the presence of Plk1 inhibitors was the cortical localization of Aur B, which was enhanced across the equator (N = 6; Figure S6C). This Aur B localization appears to concentrate at the region where the plus-ends of the astral MTs intersect. However, these cells lack Aur B concentration at a defined midzone, which is lacking in these cells. This cortical localization is intriguing, and could reflect an increase in equatorial astral MTs that deliver Aur B to the cortex – caused by the inhibition of Plk1. Alternatively, the increase in cortical Aur B could reflect an increase in the dwell time for Aur B at the cortex in response to Plk1 inhibition. This later possibility suggests the idea that Aur B normally cycles on and off of the cortex, in a Plk1-dependant manner.

Finally, we find that cells chilled in *late* anaphase in the presence of Plk1 inhibitors, then re-warmed were able to assemble symmetric furrows containing cortical Aur B (N = 3), RhoA (N = 4), and anillin (N = 4) similar to wild-type (Figure S7). We conclude that the asymmetric “half-furrows” formed when both MTs and Plk1 were inhibited in late anaphase were not due to the loss of either Plk1 or MTs alone, but instead reflect the consequence of losing both activities during late anaphase (compare Figure 3 to Figures S2 and S7).

## Conclusions

In our late-anaphase experiments, MT-cortical interactions proceed for 6 min of anaphase before chilling/drug addition, apparently a time sufficient to deliver signals necessary for furrow formation (Figure 4). If *either* MTs or Plk1 are inhibited after this point, RhoA, anillin, and Aur B all accumulate at the equatorial cortex, and ingression proceeds. This suggests that late in anaphase the equatorial cortex is “committed to furrowing”, and the RhoA/anillin scaffold no longer requires instructions from the spindle – mediated by MTs or Plk1 – in order to engage in contractility. If however, both MTs and Plk1 are inactivated after this point, then radially symmetric furrow ingression fails. Interestingly the active “half-furrow” retains the RhoA/anillin scaffold, along with Aur B, factors the inactive “half-furrow” lacks. The retention of anillin to the active furrowing surface is interesting, because this protein binds directly to the phospholipid PIP_2_ in the membrane [21, 34]. Cortical maturation, i.e. anillin anchoring at the plasma membrane, may trigger the “commitment to furrowing” stage and be driven by the increase of specific phospholipids at the furrow. This in turn would influence the retention of Rho and its effectors at the equator [19–21, 34, 36].

**Figure 4.**
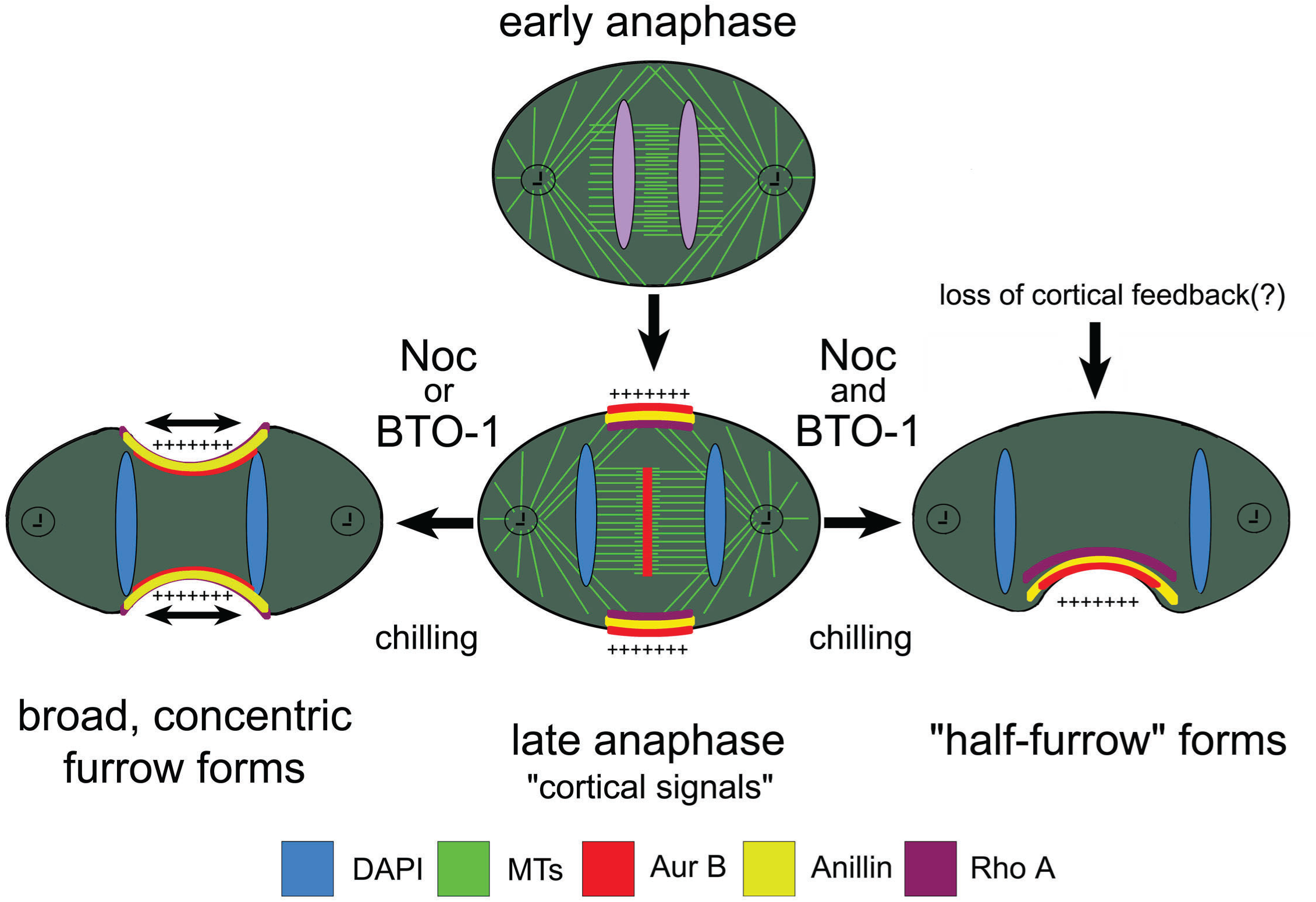
Model for “commitment to furrowing” and “half-furrow” assembly. Top cell in early anaphase, the stimulatory signals have yet to reach the equatorial cortex. As the cell progresses into late anaphase, these signals are deposited along the equatorial cortex, resulting in a prominent band of cortical Aur B kinase, anillin scaffold, and active RhoA (plus signs). At this point, removal of MTs or Plk1 kinase activity does not disrupt the retention of these factors from the cortex, and symmetric cleavage proceeds, albeit as a broadened furrow – this suggest a “commitment to furrowing”. However, removal of both MTs and Plk1 kinase activity results in the selective loss of Aur B kinase, anillin scaffold, and active RhoA from one surface of the cortex but not the other, leading to the ingression of a “half-furrow”. This suggests that by late anaphase, there are sufficient furrowing signals delivered to the equatorial cortex to support furrowing, and therefore, no further input from the spindle is required. However, this also suggests that normally cleaving cells have a mechanism that retains this furrowing activity, and such a mechanism cannot tolerate the loss of both MTs and Plk1 activity. The half-furrow represents a preferential signal, perhaps induced by acentric spindle position, which has progressed to the point of ingression, while the opposing furrow has not progressed to this point, and losses its furrow-inducing signals and structural elements.

In many cases, cleavage furrow formation is initiated asymmetrically, with only one cortical surface ingressing towards the spindle midzone – reflecting asymmetric cleavage initiation determined by relative spindle position in the cell [10–12]. In some instances, furrowing remains asymmetric, while in others the entire equatorial cortex eventually ingresses [12]. Asymmetric furrowing can also be caused by a physical block between spindle and cortex that prevents recruitment of “cleavage factors” (i.e. RhoA and anillin) to one side of the furrow, as in the case of *Chlamydia trachamatosis* infection, which induces a vacuolar inclusion in the center of a dividing cell [9]. Therefore, each hemispherical surface of the cortex can act independently, responding to temporally distinct MT-cortical interactions. Activation of furrow assembly is dependant upon the delivery of RhoA modulating enzymes to the equatorial cortex by kinesin-6 [2, 3, 18, 37, 38], and this delivery is regulated by Plk1 activity [30–32]. Our results are consistent with the idea that this delivery results in a feedback loop from the cortex/membrane that reinforces the establishment of the furrow-competent signal [39]. The Aur B deposited at the cortex by astral MTs could potentially play such a role by regulating the interactions between the RhoA/anillin scaffold and 14-3-3 proteins/actomyosin and kinesin-6 in the presumptive furrow [40, 41]. Such a feedback loop would ensure that cleavage remains robust, even in the face of asymmetric signal strength, resulting from acentric spindle localization with in the cell.

### Experimental Procedures

Unless otherwise noted all reagents were obtained from Sigma Chemical Co. (St. Louis, MO).

### Cell Culture and treatment

BSC-1 cells (African Green Monkey kidney cells; ATCC, Manassas, VA) expressing EGFP-α tubulin (Clontech, Mountain View, CA) were cultured in DMEM containing 10% FBS (Gibco, Grand Island, NY) and 1 mg/ml pen-strep (Sigma-Aldrich, St. Louis, MO) in 10% CO_2_.

MT depolymerization was done by adding Nocodazole (Sigma) to a final concentration of 33 μM. Plk1 was inhibited by the addition of 20 μM BTO-1 (EMD Millipore, Billerica, MA).

Live-cell fluorescence images were captured using a Leica DM RXA2 microscope using a Leica 63x/1.3 NA glycerol-immersion objective and enclosed in a custom-made Plexiglas box maintained at 37 °C, mounted with a Yokagawa CSU-10 spinning disk confocal head, as modified by McBain Systems (Simi Valley, CA), a Coherent 200 mW 488 nm laser (Coherent Inc. Santa Clara, CA), and a Hamamatsu 9100 back-thinned EM-CCD camera.

Cells on coverslips were either fixed in cold methanol (−20 °C), ice-cold 10% TCA, or at room temperature with 1% glutaraldehyde [23]. For glutaraldehyde fixation, cells were first extracted at the appropriate temperature for 1 minute in cytoskeleton buffer (137 mM NaCl, 5 mM KCl, 1.1 mM Na_2_HPO_4_, 0.4 mM KH_2_PO_4_, 2 mM MgCl_2_, 2 mM EGTA, 5 mM PIPES, 5.5 mM Glucose, pH 6.1) with 0.3% Triton-X 100 and 0.5% glutaraldehyde added, followed by fixation for 15 minutes in cytoskeleton buffer with 1% glutaraldehyde added. To reduce free aldehyde groups, fixation was followed by treatment with 0.5 mg/ml NaBH_4_ in cytoskeleton buffer for 5 min. Cells were labeled with combinations of the following: phalloidin-Alexa 594 (Invitrogen, Carlsbad, CA), anti-α-tubulin (mouse monoclonal D1a clone1:1000; Sigma), anti-Aurora B (1:1000, Abcam, Cambridge, MA), anti-PRC1 (1:1,000; Santa Cruz Biotechnology-SCBT, Santa Cruz, CA), anti-MyoII (1:500; Covance, Princeton, NJ), anti-MKLP1 (1:1000, SCBT), anti-RhoA, (1:100; SCBT), or anti-anillin (gift of Dr. Christine Field, Harvard Medical School). The 2° antibodies were the species appropriate Alexa-488, Alexa-594, or Alexa-660 at 1:500 dilution (Invitrogen, Carlsbad, CA). Cells were counter stained with DAPI (Sigma) and mounted in Prolong Gold with stiffener (Invitrogen, Carlsbad, CA).

For same cell live-imaging/fixed imaging, the position of the relevant cell was marked with a diamond scribe in the nosepiece of the microscope, then the coverslip is removed from the imaging chamber and fixed in the appropriate fixative. The position of the cell is relocated in the fixed cell microscope using a 10x phase contrast objective. Quantification of Aur B fluorescence intensity was done as described [42].

Fixed cells were imaged with a 63X 1.4 NA Apochromatic objective on a Leica DM RXA2 microscope using a Hamamatsu ORCA-ER CCD camera; Z-stacks were compiled as maximum projections. Images were acquired with Slidebook software (Intelligent Imaging Innovations, Denver Colorado). Figures were assembled using Photoshop 6.0 (Adobe, San Jose, CA).

### Cold-based MT depolymerization/re-polymerization assay

Cell chilling to disassemble spindle MTs during anaphase was done as described [24, 42]. Briefly, cells on coverslips assembled onto aluminum support slides were imaged by spinning disk confocal microscopy at 37 °C, and at select times, transferred to a –20 °C freezer for 5 min, followed by 15 min at 4 °C. During cell chilling, select small molecule inhibitors were added directly to the imaging chamber to their final dilution. To re-warm the cells the chambers were placed in the 37 °C microscope chamber, the cell rapidly re-focused by wide-field fluorescence microcopy, and then switched to spinning disk confocal.

## Acknowledgments

We thank Dr. Christine Field (Harvard Univ.) for the gift of anti-anillin antibody. TL and HM were each supported by the Hormel Institute’s “Summer Undergraduate Research Experience” (SURE) program. This work is supported by grants from the Department of Defense (CDMRP CA171071 and CA200747 to EHH), the Minnesota Partnership for Biotechnology and Medical Genomics (to DJD and EHH), and the National Institutes of Health (R01NS117432 to EHH and DJD). CAD is supported by the National Cancer Institute T32 CA21783602 Neuro-Oncology Training Grant to the Mayo Clinic.

## Supplemental Figures

**Supplemental Figure 1. Mitotic spindle assembly and cytokinesis following cold-based microtubule disassembly/re-warming. A.** Mitotic progression and cytokinesis in a control GFP-α tubulin expressing cell. Anaphase onset begins at T=1 min (**A**). At T=10 min (**\***) the cleavage furrow forms (arrows). Cytokinesis continues and the furrow constricts the midzone microtubules into a prominent midbody. Avg. time from anaphase onset to furrowing in these cells is 10.0 min +/- 1.6 min (N = 16). **B.** Chilling disassembles the spindle microtubules. **a.** Metaphase cell progresses into late anaphase, builds a spindle midzone, and is chilled to 4 °C (**e**). After 20 min at 4 °C, the cell is fixed. **e-e”’**. Anti-α tubulin, anti-PRC1 and DAPI reveal that the microtubules of the spindle are completely disassembled, including those of the spindle midzone. **C.** Same-cell live/fixed cell imaging of cells in late anaphase, fixed before or after cooling. The distribution of anillin (**a”**) is not affected by chilling (**b”**). The same holds true for Rho A (**c”** vs **d”**, arrows). **D.** Cold induced spindle disassembly and re-assembly. Cell enters anaphase at T = 2 min, and is then chilled, and DMSO added to the chamber. At T= 22 min (**c**), the cell is re-warmed. The spindle re-appears and a furrow forms and constricts the cell (**f**). **E.** Avg. time from re-warming to furrow induction for cells chilled in early, mid, and late anaphase. Hrs:Min. Bar = 10 μm.

**Supplemental Figure 2. RhoA and anillin are prevented from localizing to the presumptive furrow in cells chilled with Noc during early, but not late anaphase. A.** Rho A does not localize to the cell cortex in cells treated with Noc in early anaphase (**d”**). **B.** Cells lacking cortical contraction do not accumulate anillin (**d”**). **C-D.** Same cell live-fixed images of GFP α tubulin-expressing cells chilled in late anaphase and treated with Noc. These cells assemble a cleavage furrow, which initiates asymmetrically from one surface. Both Rho A (**C**) and anillin (**D**) localize along the equatorial cortex. Bar = 10 μm.

**Supplemental Figure 3. Cortical contractility and displacement of furrowing components actomyosin, anillin, and RhoA induced by MT depolymerization during early anaphase. A.** Control cell, showing distribution of MTs (**a**), MyoII (**b**), F-actin (**c**) to the cleavage furrow. **B-D.** Same cell live-fixed series of GFP α tubulin-expressing cells, chilled in early anaphase, and undergoing uncoordinated cortical contractility. The equatorial cortical region of these cells lacks concentrations of Myosin II and F-actin (**A**), anillin (**B**), or RhoA (**C**). Bar = 10 μm.

**Supplemental Figure 4. Aur B redistribution to the equatorial cortex occurs late in anaphase and requires the astral microtubules. A.** Cell chilled in early anaphase (**c**), then re-warmed in the presence of depolymerizing dose of Nocodazole (33 μM). The spindle does not reform, and Aur B remains associated with the chromosomes, but does not localize to the equatorial cortex (**e”**). **B.** Cell chilled in early anaphase (**b**), then re-warmed in the presence of a non-depolymerizing dose of Nocodazole (0.25 μM). The dynamic asters do not reform, but there are midzone MTs. In these cells the chromosomal Aur B re-distributes to the midzone; however, in these anastral cells Aur B fails to localize to the equatorial cortex (**f”**, arrows). Compare Aur B localization to nuclei from top and bottom panels (**e”’** vs. **f’”**). Hrs:Min. Bar = 10 μm.

**Supplemental Figure 5. Inhibiting Plk1 activity during re-warming prevents cortical contractility and furrowing. A.** BTO-1-treated cell following chilling-re-warming. Cell in early anaphase (2 min; **c**), then was chilled, and BTO-1 added. As the cell was re-warmed extensive astral MTs re-form, but the cell does not undergo cortical contractility and does not form a furrow (**e**). Aur B does not re-distribute from the chromosomes, and there is not equatorial cortical localization of Aur B **B.** BTO-1 treated cell. This cell progressed into early anaphase (2 min; **c**), and then was chilled and BTO-1 was added. Once re-warmed, the astral MT arrays reform, but not midzone assembles, and there is no cortical contractility. Note that the chromosomal Aur B is lost, and there is extensive localization of Aur B to the equatorial cortex. **C.** Average time from re-warming to the induction of a furrow for cells chilled and treated with BTO-1 inhibitors in early, mid, and late anaphase. Time is hrs:min. Bar = 10 μm.

**Supplemental Figure 6. Cells fail to reform a functional midzone MT array when Plk1 is inhibited during early anaphase.** Three cells allowed to progress for 2 min into anaphase, then chilled and treated with BTO-1. Following re-warming, these cells were fixed and immuno-labeled with antibodies to two separate central spindle components: **A.** MKLP1, and **B.** PRC1. **A.** The cell has reformed astral MT arrays, but lacks an MKLP1-positive spindle midzone. Note the MKLP1-positive midbody on the adjacent cell (arrow in **e”-e’”**). **B.** Post-warming cell lacks a PRC1 positive midzone MT. Identical results were obtained when Plk1 activity was inhibited with BTO-1. Cells do not initiate cortical contractility, and lack MKLP1-positve central spindle (**C**) or PRC1 (**D**). Time in lower right hrs:min. Bar = 10 μm.

**Supplemental Figure 7. Loss of Plk1 activity in late anaphase cells does not prevent furrow formation.** Three cells allowed to progress for 6 min into anaphase, then chilled and treated with BTO-1. The cells were re-warmed after 20 min. After the spindle MTs reformed the cells were fixed and labeled with antibodies against Aur B (**A**), anillin (**B**) or Rho A (**C**). Blocking Plk1 activity in late anaphase does not inhibit the ability of Aur B, anillin, or Rho A from localizing to the furrow (arrows in g”), nor does it prevent circumferential furrowing. Time in lower right hrs:min. Bar = 10 μm.

## References

1. D’Avino, P.P, M.S. Savoian, and D.M. Glover (2005). Cleavage furrow formation and ingression during animal cytokinesis: a microtubule legacy. J. Cell Sci. 118:1549–1558.

2. von Dassow, G. (2009). Concurrent cues for cytokinetic furrow induction in animal cells. Trends Cell Biol. 19:165–172.

3. Bement, W.M., H.A. Benink, and G. von Dassow (2005). A microtubule-dependent zone of active RhoA during cleavage plane specification. J. Cell Biol. 170:91–101.

4. Yüce, O., A. Piekny, and M. Glotzer (2005). An Ect2-centralspindlin complex regulates the localization and function of RhoA. J Cell Biol. 170:571–582.

5. Piekny, A.J., and M. Glotzer (2008). Anillin is a scaffold protein that links RhoA, actin, and myosin during cytokinesis. Curr. Biol. 18:30–36.

6. Hickson, G.R.X., and P.H. O’Farrell (2008). Rho-dependent control of anillin behavior during cytokinesis. J. Cell Biol. 180:285–294.

7. Rappaport, R. (1985). Repeated furrow formation from a single mitotic apparatus in cylindrical sand dollar eggs. J. Exp. Zool. 234:167–171.

8. Rappaport, R., and B. Rappaport (1983). Cytokinesis: Effects of blocks between the mitotic apparatus and the surface of furrow establishment in flattened echinoderm eggs. J. Exp. Zool. 227:213–227.

9. Sun, H.S., A. Wilde, and R. Harrison (2011). *Chlamydia trachomatis* inclusions induce asymmetric cleavage furrow formation and ingression failure in host cells. Mol. Cell Biol. 31:5011–5022.

10. Sanger, J.M., B. Mittal, J.S. Dome, and J.W. Sanger (1989). Analysis of cell division using fluorescently labeled actin and myosin in living PtK2 cells. Cell Motil. Cyto. 14:201–219.

11. Shuster, C.B. and D.R. Burgess (2002). Transitions regulating the timing of cytokinesis in embryonic cells. Curr. Biol. 12:854–858.

12. Maddox, A.S., L. Lewellyn, A. Desai, and K. Oegema (2007). Anillin and the septins promote asymmetric ingression of the cytokinesis furrow. Dev. Cell. 12:827–835.

13. Canman, J.C, D.B. Hoffman, and E.D. Salmon (2000). The role of pre-and post-anaphase microtubules in the cytokinesis phase of the cell cycle. Curr. Biol. 10:611–614.

14. Alsop, G.B., and D. Zhang (2003). Microtubules are the only structural constituent of the spindle apparatus required for induction of cell cleavage. J Cell Biol. 162:383–390.

15. Strickland, L.I., E.J. Donnelly, and D.R. Burgess (2005). Interaction between EB1 and p150 ^Glued^ is required for anaphase astral microtubule elongation and stimulation of cytokinesis. Curr. Biol. 15:2249–2255.

16. Murthy, K., and P. Wadsworth (2008). Dual role for microtubules in regulating cortical contractility during cytokinesis. J. Cell Sci. 121:2350–2359.

17. Foe, V.E., and G. von Dassow (2008). Stable and dynamic microtubules coordinately shape the myosin activation zone during cytokinetic furrow formation. J. Cell Biol. 183:457–470.

18. Vale, R.D., J.A. Spudich, and E.R. Griffis (2010). Dynamics of myosin, microtubules, and Kinesin-6 at the cortex during cytokinesis in *Drosophila* S2 cells. J. Cell Biol. 186:727–738.

19. Gregory, S.L., S. Ebrahimi, J. Milverton, W.M. Jones, A. Bejsovec, and R. Saint (2008). Cell division requires a direct link between microtubule-bound RacGAP and anillin in the contractile ring. Curr. Biol. 18:25–29.

20. Kechad, A., S. Jananji, Y. Ruella, and G.R.X. Hickson (2012). Anillin acts as a bifunctional linker coordinating midbody ring biogenesis during cytokinesis. Curr. Biol. 22:197–203.

21. Liu, J., G.D. Fairn, D.F. Ceccarelli, F. Sicheri, and A. Wilde (2012). Cleavage furrow organization requires PIP_2_-mediated recruitment of anillin. Curr. Biol. 22:64–69.

22. Murata-Hori, M., and Y-l. Wang (2002). Both midzone and astral microtubules are involved in the delivery of cytokinesis signals: insights from the mobility of aurora B. J. Cell Biol. 159:45–53.

23. Hornick, J.E., C.C. Mader, E.K. Tribble, C.C. Bagne, K.T. Vaughan, S.L. Shaw, and E.H. Hinchcliffe (2011). Amphiastral mitotic spindle assembly in vertebrate cells lacking centrosomes. Curr. Biol. 21:598–605.

24. Bader, J.R., J. Kasuboski, M. Winding, C. Zhang, P.S. Vaughan, E.H. Hinchcliffe, and K.T. Vaughan (2011). Polo-like kinase 1 is required for recruitment of dynein to kinetochores during mitosis. J. Biol. Chem. 286:20769–20777.

25. Mollinari, C., J-P. Kleman, Y. Saoudi, S.A. Jablonski, J. Perard, T.J. Yen, and R.L. Margolis (2005). Ablation of PRC1 by small interfering RNA demonstrates that cytokinetic abscission requires a central spindle bundle in mammalian cells, whereas completion of furrowing does not. Mol. Biol. Cell. 16:1043–1055.

26. Durcan, T., E.S. Halpin, T. Rao, N. Collins, E.K. Tribble, J.E. Hornick, and E.H. Hinchcliffe (2008). Tektin 2 is required for central spindle microtubule organization and the completion of cytokinesis. J. Cell Biol. 181:595–603.

27. Hornick, J.E., K.B. Karanjeet, E.S. Collins, and E.H. Hinchcliffe (2010). Kinesins to the core: The role of microtubule-based motor proteins in building the mitotic spindle midzone. Semin. Cell Dev. Biol. 21:290–299.

28. Bringmann, H., and A.A. Hyman (2005). A cytokinesis furrow is positioned by two consecutive signals. Nature. 436:731–734.

29. Miller, A.L., and W.M. Bement (2009). Regulation of cytokinesis by Rho GTPase flux. Nat. Cell Biol. 11:71–77.

30. Brennan, IM, Peters U, Kapoor TM, Straight AF (2007) Polo-like kinase controls vertebrate spindle elongation and cytokinesis. PLoS ONE 2(5):e409. doi:10.1371/journal.pone.0000409

31. Petronczki, M., M. Glotzer, N. Kraut, and J-M. Peters (2007). Polo-like kinase 1 triggers the initiation of cytokinesis in human cells by promoting recruitment of the RhoGEF Ect2 to the central spindle. Dev. Cell 12:713–725.

32. Burkhard, M.E., C.L. Randall, S. Larochelle, C. Zhang, K.M. Shokat, R.P. Fisher, and P.V. Jallepalli (2007). Chemical genetics reveals the requirement for Polo-like kinase 1 activity in positioning RhoA and triggering cytokinesis in human cells. PNAS 104:4383–4388.

33. Wang, Y-l. (2005). The mechanism of cortical ingression during early cytokinesis: thinking beyond the contractile ring hypothesis. Trends in Cell Biol. 15:581–588.

34. Brill, J.A. R. Wong, and A. Wilde (2011). Phosphoinositide function in cytokinesis. Curr. Biol. 21:R930–R934.

35. Piekny, A.J., and A.S. Maddox (2010). The myriad roles of anillin during cytokinesis *Semin*. Cell Dev. Biol. 21:881–891.

36. Su, K-C., T. Takaki, and M. Petronczki (2011). Targeting of the RhoGEF Ect2 to equatorial membrane controls cleavage furrow formation during cytokinesis. Dev. Cell 21:1104–1115.

37. Lekomtsev, S., K-C. Su, V.E. Pye, K. Blight, S. Sundaramoorthy, T. Takaki, L.M. Collinson, P. Cherepanov, N. Divecha, and M. Petronczki (2012). Centralspindlin links the mitotic spindle to the plasma membrane during cytokinesis. Nature 492:276–279.

38. Lewellyn, L., A. Carvalho, A. Desai, A. Maddox, and K. Oegema (2011). The chromosomal passenger complex and centralspindlin independently contribute to contractile ring assembly. J. Cell Biol. 193:155–169.

39. Surcel, A., Y-S. Kee, T. Luo, and D.N. Robinson (2010). Cytokinesis through biochemical-mechanical feedback loops. Semin. Cell Dev. Biol. 21:866–873.

40. Douglas, M. E., T. Davies, N. Joseph, and M. Mishima (2010). Aurora B and 14-3-3 coordinately regulate clustering of centralspindlin during cytokinesis. Curr. Biol. 20: 927–933.

41. Zhou, Q., Y.-S. Kee, C.C. Poirier, C. Jelinek, J. Osborne, S. Divi, A. Surcel, M.E. Will, U.S. Eggert, A. Müller-Taubenberger, P.A. Iglesias, R.J. Cotter, and D.N. Robinson (2010). 14-3-3 coordinates microtubules, Rac, and Myosin II to control cell mechanics and cytokinesis. Curr. Biol. 20:1881–1889.

42. Kasuboski, J.M., J.R. Bader, S.B.F. Tauhata, P.S. Vaughan, M. Winding, M.M. Morrissey, M.V. Joyce, W. Boggess, L. Vos, G.K. Chan, E.H. Hinchcliffe, and K.T. Vaughan (2011). Zwint-1 is a novel Aurora B substrate required for the assembly of a dynein-binding platform on kinetochores. Mol. Biol. Cell 22:3318–3330.

